# Distinct changes in morphometric networks in aging versus Alzheimer’s disease dementia

**DOI:** 10.1101/615401

**Authors:** Alexa Pichet Binette, Julie Gonneaud, Jacob W. Vogel, Renaud La Joie, Pedro Rosa-Neto, D. Louis Collins, Judes Poirier, John C.S. Breitner, Sylvia Villeneuve, Etienne Vachon-Presseau, Alzheimer’s Disease Neuroimaging Initiative, PREVENT-AD Research Group

**Author notes:** Senior Author (Sylvia Villeneuve, Etienne Vachon-Presseau). Data used in preparation of this article were obtained from the Alzheimer’s Disease Neuroimaging Initiative (ADNI) database. As such, the investigators within the ADNI contributed to the design and implementation of ADNI and/or provided data but did not participate in analysis or writing of this report. A complete listing of ADNI investigators can be found here.

## Abstract

Brain gray matter (GM) morphometric changes are prevalent in both aging and Alzheimers disease (AD), though disentangling these two processes has proved challenging. Using independent component analysis, we derived morphometric networks from a large, multi-cohort dataset, and investigated how GM volume within these networks differs in young adulthood, old adulthood, and AD. Aging and AD contributed additive effects on GM loss in nearly all networks, except frontal lobe networks, where GM reductions were more specific to aging. While no networks show GM loss highly specific to AD, a higher degree of variability in the whole-brain pattern of GM volume characterized AD only. Preservation of the whole-brain GM pattern in cognitively normal older adults was related to better cognition and lower risk of developing cognitive impairment. These results suggest both aging and AD involve widespread atrophy, but that cognitive impairment is uniquely associated with disruption of morphometric organization.

## 1. Introduction

Alzheimer’s disease (AD) and normal aging are both characterized by considerable atrophy. Because age is the main risk factor for AD (Association, 2017), these two processes may be closely intertwined. Disentangling brain changes specific to aging versus AD has been a challenge (Fjell et al., 2014; Jagust, 2013). For example, whether AD neurodegeneration represents accelerated aging or a distinct process has not been fully resolved (Brayne and Calloway, 1988; Buckner, 2004; Ghosh et al., 2011; Toepper, 2017). We sought further insight into this topic by examining grey matter (GM) changes across the lifespan and AD conjointly.

AD brings neurodegeneration in several regions, especially the hippocampus, the temporal lobe and associative areas (Bakkour et al., 2013; Besson et al., 2015; Du et al., 2001; Jack Jr et al., 2015a; Wirth et al., 2013). In aging, GM atrophy in the frontal lobe is consistently reported as a principal contributor to age-related cognitive changes (Fjell and Walhovd, 2010; Resnick et al., 2003), but the temporal lobe seems also particularly vulnerable to advancing age, even in elderly at low risk of AD (Fjell et al., 2013). While studies investigating large-scale structural networks are less numerous, the pattern of atrophy in AD dementia seems to mimic functional and GM covariance networks (Seeley et al., 2009). GM covariance networks may also change with advancing age (DuPre and Spreng, 2017; Koini et al., 2018), and possibly more so in AD relative to aging (Spreng and Turner, 2013). Together, these findings suggest an additive effect of aging and AD on volume change in certain brain regions and/or on the whole-brain structural organization. This raises questions as to which GM changes, if any, are specific to aging or AD (Jagust, 2013). Discerning features specific to AD beyond those of aging could suggest novel ways to consider neurodegeneration in the AD research framework.

We applied independent component analysis (ICA) to GM maps from individual structural MRI of participants from a large, multi-cohort dataset spanning young adults, older adults with intact cognition and with AD dementia, to derive morphometric networks, a term used as an analogy to functional networks created by ICA of functional MRI data. We investigated GM volume changes within these morphometric networks, along with changes in their intrinsic organization. Our analyses were framed around a hypothetical model that relegated GM changes between groups into three classes, one being disease-specific (Figure 1A), one being characteristic of aging alone (Figure 1B), and one representing an additive effect of both (Figure 1C).

**Figure 1:**
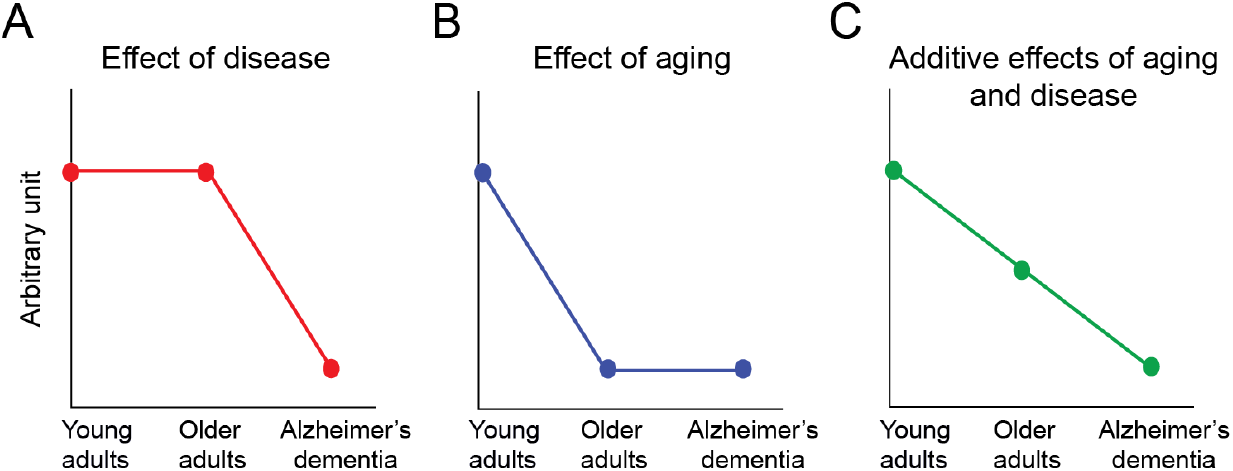
Three proposed trajectories of grey matter (GM) changes between groups. A) Effect of disease: a GM feature similar between young and older adults, but altered in AD. B) Effect of aging: a GM feature similar between older adults with and without AD, but different compared to young adults. C) Additive effect of aging and disease: a GM feature changing gradually across lifespan and AD continuum. The y-axis represents the magnitude of change in morphometric networks and/or intrinsic organization. The x-axis represents different conditions.

In this study, we first uncovered data-driven morphometric networks that were stable across all individuals using ICA. Age had an impact on all networks, and GM volume loss in most networks showed an additive effect of age and AD. The inter-individual variability of GM volume across networks was similar in young and cognitively normal older adults. AD was specifically characterized by higher variability across and between morphometric networks, resulting in a whole-brain pattern that was more heterogeneous than what was found in young adults and in normal aging. Furthermore, having a whole-brain pattern less similar to young adults was associated with worse cognition and increased risk of developing cognitive impairment. These findings suggest that as long as whole-brain GM organization is preserved, individuals can remain cognitively normal, even if they have severe atrophy.

## 2. Results

### 2.1. Deriving morphometric networks

Different cohorts of young adults, older adults with intact cognition, and along the AD clinical continuum (n=1019, Table 1) were processed under a unified pipeline in which each participant’s GM map was registered to a common template. The resulting 1019 GM maps were used as input for an ICA to derive 30 principle components, which explained 62% of variance in the data. The principle components were thresholded and binarized to retain the most significant voxels and are hereafter referred to as morphometric networks. The 30 morphometric networks are shown in Figure 2A and their anatomical description can be found in Table S1. Most morphometric networks were reminiscent of clearly defined anatomical regions, such as the precuneus, basal ganglia, occipital cortex or the thalamus. All networks showed a bilateral distribution, except network 23 and 26 that encompassed the part of the left occipital lobe and the right temporal lobe, respectively. The average GM volume was extracted from each of the 30 morphometric networks, and these values formed the basis of all subsequent analyses.

**Figure 2:**
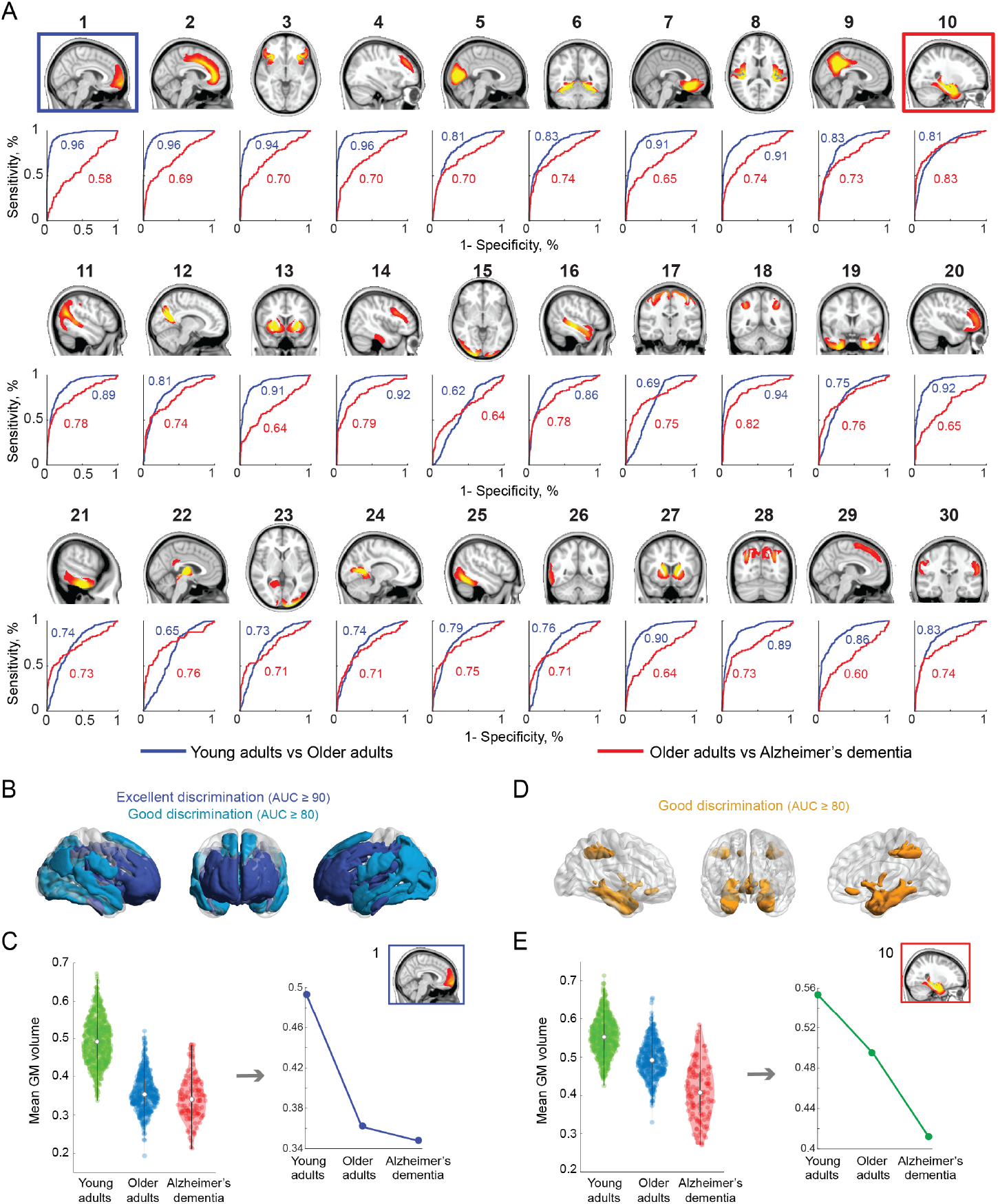
Performance of each morphometric network to discriminate aging and AD. A) The 30 anatomically derived morphometric networks from the ICA thresholded at Z ≥ 3.5. Ten-fold cross-validation was used to determine the performance of each network to discriminate between Young and Older adults (blue ROC curves) and Older adults and Alzheimer’s dementia (red ROC curves). The blue square highlights the most discriminative network for normal aging and the red square highlights the most discriminative network for Alzheimer’s dementia. B) Networks with excellent (AUC ≥ 90) and good (AUC ≥ 80) accuracy to discriminate normal aging. C). Average GM volume in the best age-related

**Table 1:**
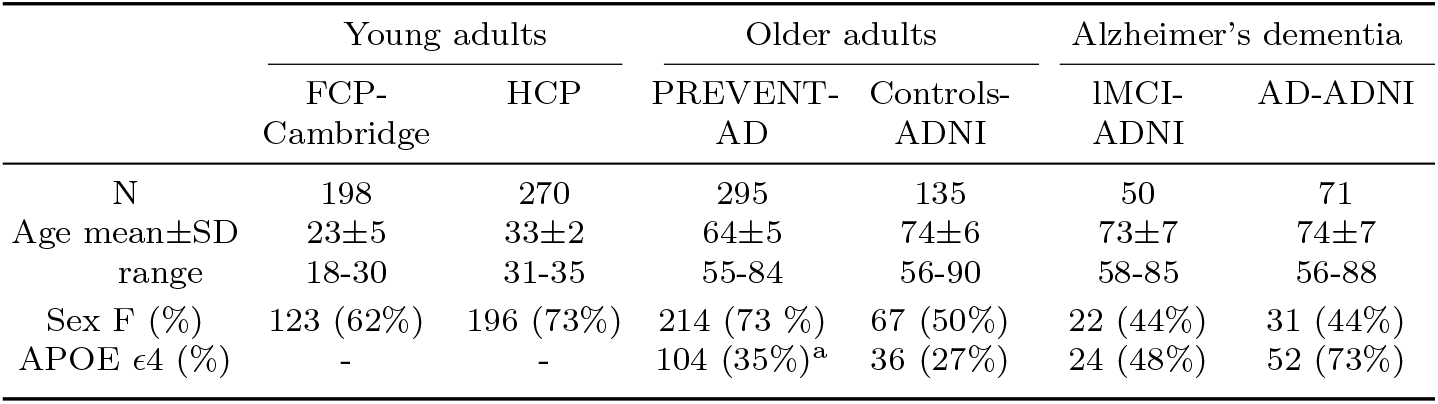
Demographics. Individuals were classified as APOE4 carriers if at least one allele is *∊*4. ^a^APOE status was available for 287 PREVENT-AD participants APOE=apoliprotein; FCP-Cambridge=1000 Functional Connectomes project - Cambridge site; HCP=Human Connectome Project; lMCI=late mild cognitive impairment; AD=Alzheimer’s disease; ADNI=Alzheimer’s disease neuroimaging initiative; SD=standard deviation.

To evaluate whether the morphometric networks would be biased by patients with severe cognitive impairment, the same ICA approach was applied to lMCI and AD participants only. Qualitatively, similar morphometric networks were identified between those two groups and all participants Figure S1.

### 2.2. Additive effect of age and AD on GM volume was found in most morphometric networks

The GM volume across morphometric networks differed between cohorts (see repeated measures ANOVA in Figure S2), showing effects of age and disease. To diminish potential confound of site effects, we combined the six cohorts into three groups: “Young adults” (FCP-Cambridge and HCP), “Older adults” (PREVENT-AD and Controls-ADNI) and “Alzheimer’s dementia” (late mild cognitive impairment [lMCI]- and AD-ADNI), and examined the general differences between these three groups.

We used a ten-fold cross-validated logistic regression procedure to determine if the GM volume in each of these morphometric networks could classify Young adults vs. Older adults and Older adults vs. Alzheimer’s dementia in the left out subjects. The AUCs from the ROC analyses represent the overall performance of each morphometric network to classify participants across the collated test sets (Figure 2A).

Many of the AUCs showed excellent (AUCs ≥ 90, n=11) or good (80 ≤ AUCs < 90, n=10) performance for classifying Young vs. Older adults (Figure 2B). Only three networks including the motor cortex (network 15), the visual cortex (network 17) and the thalamus/brain stem (network 22) performed poorly (AUCs ≤ 69). The medial prefrontal cortex (network 1, Figure 2C) was best at discriminating Young from Older adults (AUC=0.96) and could not discriminate Older adults from Alzheimer’s dementia (AUC=0.58). GM decreased from youth to old age in this network, but was stable from older adulthood to dementia - suggesting that this network is more specific to aging than to AD (Figure 2C). The AUCs of the classifiers stratifying Older adults vs. Alzheimer’s dementia were lower, with no AUC being excellent and only two being good discriminators (Figure 2D). The medial temporal network including the hippocampus and amygdala (network 10, Figure 2E) best discriminated Older adults from Alzheimer’s dementia (AUC=0.83). Interestingly, the second best network to discriminate Older adults and Alzheimer’s dementia (network 18) included part of the supramarginal and angular gyri, brain regions that have been shown repetitively to be affected by AD (Dickerson et al., 2011; Landau et al., 2011). However, GM volume in these networks (Figure 2E showing network 10), as in most other networks, showed an additive effect of age and disease.

### 2.3. Disruption of intrinsic whole-brain GM pattern in AD

GM volume signatures across morphometric networks for each participant are shown in Figure 3A. Based on those values, we derived metrics reflecting whole-brain GM pattern similarity by correlating the GM volumes signatures of the 30 morphometric networks between every other participant (Figure 3B shows a signature for two participants). This multivariate analysis captured the variability of individuals with their own group as well as with other groups. We averaged the subject-to-subject GM signature correlations for each pair-wise group, as a measure of the intrinsic GM pattern within-group (diagonal elements of matrix 3C), which ranged from 0.64 to 0.82. The intrinsic GM patterns within the groups of Young and within the groups of Older adults were homogeneous, while the pattern was less organized in AD with lower mean correlation values (Figure 3C,D) and higher standard deviation (Figure 3E). At the individual level, intrinsic GM pattern measure (within-group correlation) discriminated Older adults vs. Alzheimer’s dementia (AUC=0.72), but not Young vs. Older adults (AUC=0.57; Figure 3F).

**Figure 3:**
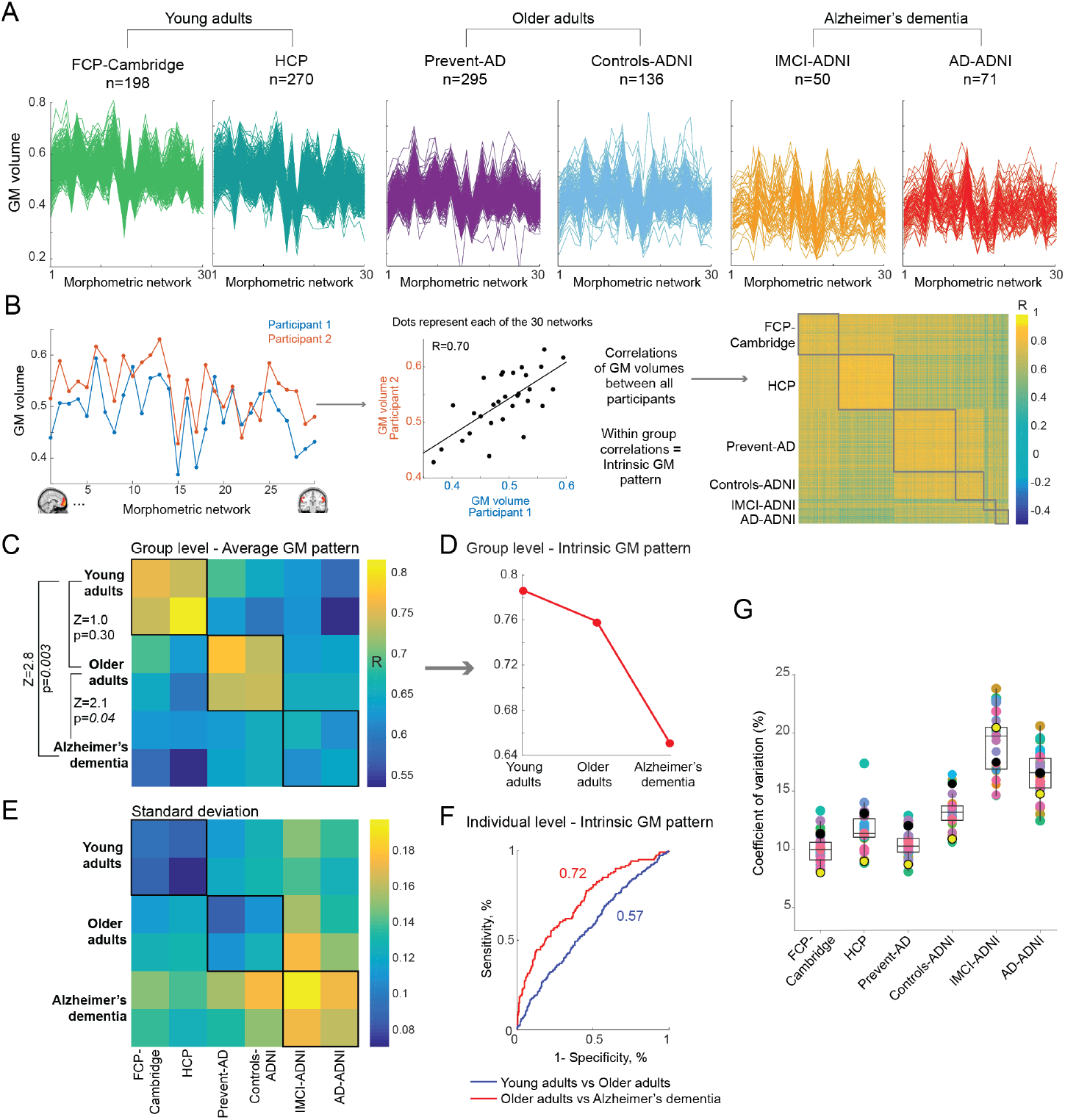
Intrinsic grey matter pattern discriminated between aging and Alzheimer’s disease. A) GM volume (y-axis) across the 30 morphometric networks (x-axis) for all participants in each of the six groups. All axes are on the same scale. B) Measures of GM pattern were derived by correlating the GM volumes across the 30 networks of each participant to every other participant. This resulted in a matrix of 1019×1019, comparing the GM pattern between all subjects. C) The average correlation of GM pattern between and within (diagonal elements) groups. Statistical differences between the intrinsic GM pattern (within group correlations) in Young adults, Older adults, and Alzheimer’s dementia are reported on the left of the matrix. D) Intrinsic GM pattern is preserved in aging, but not in AD, following the disease model. E) Standard deviations of GM pattern between and within groups. F) ROC curves showing the discriminative accuracy between Young and Older adults and between Older adults and Alzheimer’s dementia based on individual measures of intrinsic GM pattern in a ten-fold cross-validation procedure. G) Coefficients of variation (standard deviation / mean) of GM volume in the 30 networks across groups. Each dot represents a brain network. Black dots correspond to the age-related network (network 1) and yellow dots, to the AD-related network (network 10). See also Figure S3.

Although Young and Older adults showed a coherent pattern within their respective groups, the pattern itself, however, changed with aging and with AD (off-diagonal elements Figure 3C). Figure S3 shows that the GM signature correlation values can differentiate between Young and Older adults (AUC=0.94) and between Older adults and AD dementia (AUC=0.85). Our results therefore suggest that GM changes happen in a coherent way across networks in normal aging, but not in AD. Thus, higher heterogeneity and a disrupted whole-brain pattern are specific characteristics of AD, in line with the disease model (Figure 1A).

### 2.4. GM volume heterogeneity is higher in AD but not in normal aging

In line with the loss of GM pattern organization in AD, there was higher heterogeneity of GM volumes across morphometric networks in AD, as assessed by comparing coefficients of variation of GM volume. There was a main effect of group on coefficients of variation on the 30 networks (all modified signed-likelihood ratio [MSLR] tests > 33.4, p-values < 0.001). Young and Older adults showed lower variation (mean coefficient of variation in the 30 networks of 10.8 and 11.8% respectively), while Alzheimer’s dementia groups showed higher het-erogeneity (mean coefficient of variation of 17.8%) (Figure 3G). The absence of higher heterogeneity over the course of aging was validated using the Cambridge Centre for Ageing and Neuroscience (Cam-CAN) study, a mono-centric lifespan study (n=647; age range 18 to 88 years old, Figure 4A). The coefficients of variation of GM volume in the 30 morphometric networks projected on the Cam-CAN maps were similar across decades in 26 networks (all p-values > 0.004 from MSLR tests; mean coefficient of variation across decades ranged from 10.5 to 14.1%; Figure 4B). Such results challenge the proposition that normal aging significantly amplifies heterogeneity of GM volume. Instead, our results suggest that higher inter-individual variability in GM volume may be a hallmark of AD.

**Figure 4:**
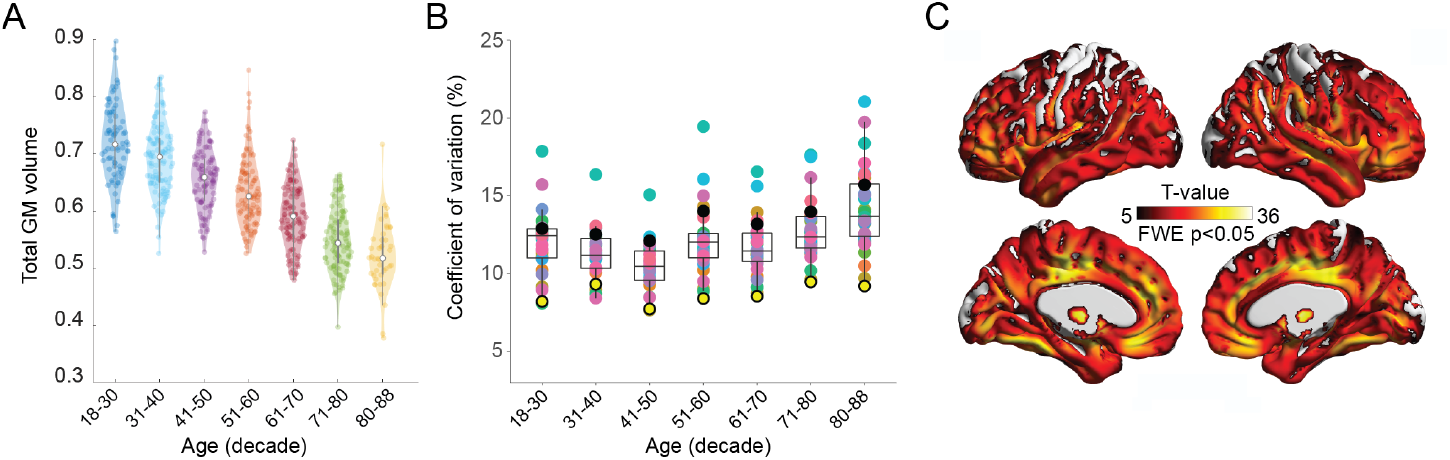
Results related to aging were validated using the Cam-CAN dataset. A) Reduction in GM total volume with advancing age. B. Similar variability of GM volume in the 30 morphometric networks across decades. Each dot represents a brain system. Black dots correspond to the age-related network (network 1) and yellow dots, to the AD-related network (network 10). C) Voxel wise analysis showed that the peaks of GM volume reduction associated with age were located in the medial prefrontal cortex, the dorsolateral prefrontal cortex, the cingulate cortex and the medial temporal lobe. Statistical significance is set at p < 0.05 family-wise error (FWE) corrected. See also Table S2.

A voxel-wise analyses of age confirmed a whole brain reduction of GM volume (Figure 4C). Not surprisingly, the peaks showing the strongest relationship with advancing aging were located in the morphometric networks with the highest accuracy to discriminate Young from Older adults.

### 2.5. Cognitive performance and clinical progression are related to a preserved GM pattern

Finally, we evaluated the clinical validity of different GM features by assessing whether they were related to cognitive performance or clinical progression in cognitively normal older adults. We focused on GM volume in the most discriminative morphometric network between Young and Older adults (age-related network, network 1) and the most discriminative between Older adults and Alzheimer’s dementia (AD-related network, network 10), along with a metric of preserved whole-brain pattern (similarity to young adults, i.e. correlation between GM volume in theF p 30 networks to the mean GM volumes of Young adults in the 30 networks).

Looking at cognitive performance in PREVENT-AD, we found that participants with a GM pattern more similar to young adults had better executive function and a trend toward better memory performance (Table 2). There was no association between cognition and GM volume in the age-related network, but lower GM volume in the AD-related network was associated to worse memory performance. Performing similar analyses in Controls-ADNI, a cohort on average ten years older than PREVENT-AD, revealed consistent findings; participants showing a GM pattern more similar to young adults had better cognitive performance.

**Table 2:** Relationships between cognitive performance and GM features in cognitively normal older adults. Results from separate linear regression models showing how GM pattern similarity to the young adults and GM volume in the age and AD systems (independent variables) are related to cognitive performance. Models included education and total GM as covariates. p-values are not corrected for multiple comparisons.

|  | GM pattern |  | GM volume |  |  |  |
| --- | --- | --- | --- | --- | --- | --- |
|  | Similarity to<br>young adults |  | Age-related<br>morphometric<br>network 1 |  | AD-related<br>morphometric<br>network 10 |  |
|  | F | p | F | p | F | p |
| **PREVENT-AD** (n=291) |  |  |  |  |  |  |
| List learning (memory) | 3.50 | 0.06 | 0.95 | 0.33 | 9.02 | <0.01 |
| Coding (executive function) | 5.85 | 0.02 | 0.01 | 0.76 | 0.15 | 0.70 |
| **Controls-ADNI** (n=135) |  |  |  |  |  |  |
| ADAS-Cog | 8.09 | <0.01 | 1.65 | 0.20 | 0.03 | 0.85 |

In Controls-ADNI, a proportion of participants converted to MCI (n=18), most of them between 2 to 4 years later. When compared to Controls-ADNI who remained cognitively normal (n=117), these converters displayed a GM pattern less similar to young adults (Figure 5A). Trends toward lower GM volume in the age- and the AD-related networks were found in converters when compared to stable older adults (Figure 5B,C). Using leave-one-out cross-validation analyses, we showed that whole-brain pattern similarity to Young adults differentiated Controls-ADNI converters from stable with a fair accuracy (AUC=0.71), whereas GM volume in the age- and AD-related networks yielded poor accuracy (Figure 5, bottom row). These findings, consistent across two independent cohorts of cognitively normal older adults, support the previous results suggesting that whole-brain GM organization is an important feature of clinical manifestation of cognitive impairment.

**Figure 5:**
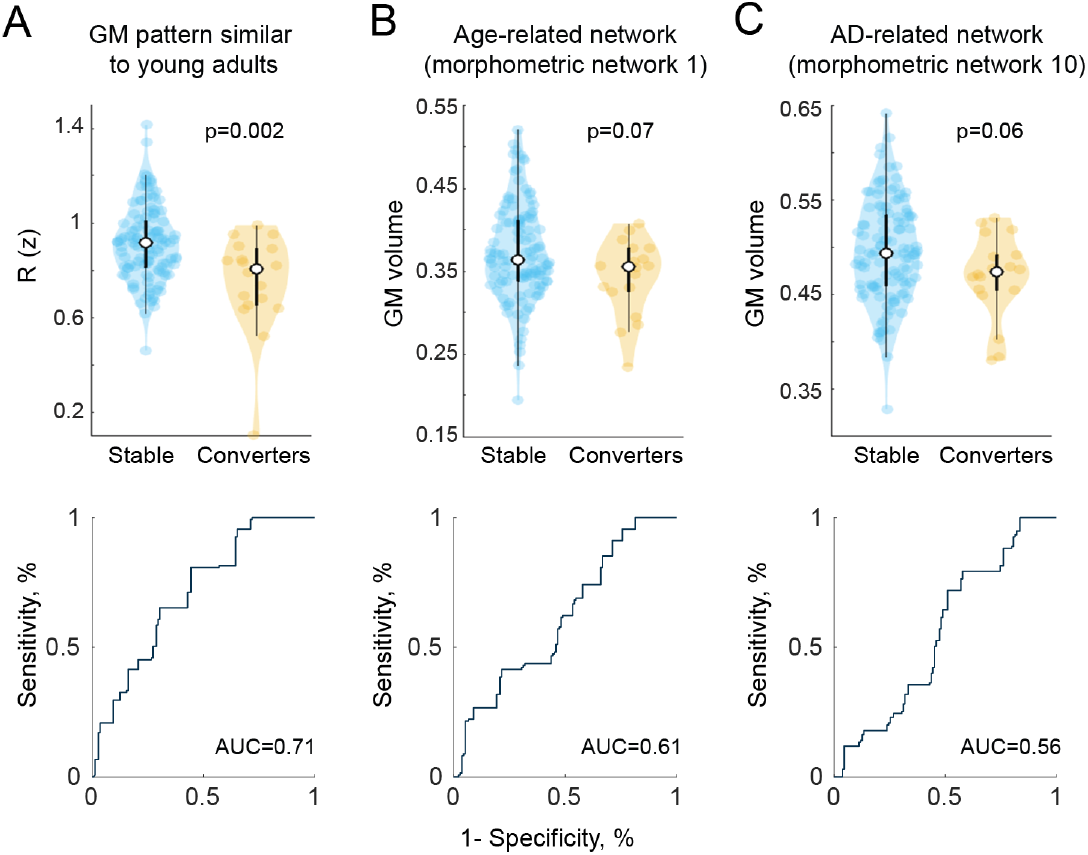
Whole-brain GM organization related with cognitive decline. Differences between Controls-ADNI who converted to MCI (Converters, n=18) and those who remained cognitively normal (Stable, n=117) on GM pattern similarity to young adults (A), GM volume in the age-related network (B) and the AD-related network (C). P-values from Mann-Whitney U tests and not corrected for multiple comparisons. Bottom row shows ROC curves to discriminate between Stable and Converters. Results remained the same when excluding one extreme case with the lowest GM pattern similarity

## 3. Discussion

Using a large, multi-cohort dataset, we identified a set of 30 morphometric networks, and evaluated how GM volume changes in these networks, individually and in concert, during the course of aging and AD. We used cross-validation procedures to determine how each feature could discriminate young from cognitively normal older adults (effect of age) and cognitively normal older adults from Alzheimer disease (effect of the disease). Across the whole brain, we observed an important decrease in GM volume in the course of aging, as almost all morphometric networks could accurately stratify young adults from older adults. Atrophy related to AD added to that of aging in most brain systems, excluding those in the medial frontal cortex. Importantly, AD, but not aging, was associated with increased heterogeneity in GM volume across the morphometric networks and in whole-brain GM pattern. The robustness of the results was validated in the Cam-CAN monocentric lifespan cohort, where GM volume variability was consistent across the decades. Finally, having a GM pattern less similar to young adults was related to progression to MCI in Controls-ADNI.

How does the brain age? Is AD a form of accelerated aging? What features distinguish changes of normal aging from those seen in early AD? To disentangle changes of normal aging vs. those leading to neurodegenerative diseases, large longitudinal studies monitoring structural and pathological brain changes across lifespan would be needed. While such studies do not exist, several lifespan and disease cohorts are now available, making it possible to infer longitudinal changes based on large cross-generational data. Using more than a thousand structural MRI scans from adults aged 18 to 89 years old, among which 12% were diagnosed with lMCI or AD dementia, we differentiated brain changes more specific to AD from those more specific to aging and identified those vulnerable to both phenomena. We were interested in both the magnitude (volume) and the pattern (whole-brain organization) of GM features. Also, rather than targeting a priori structural brain regions, we used ICA to uncover 30 morphometric networks that were representative of our sample and therefore not biased by one specific group of interest (Bassett et al., 2008; Hafkemeijer et al., 2014; Zeighami et al., 2015).

Because frontal systems are preferentially affected by age but not by AD, our results do not support the hypothesis that AD-related neurodegeneration simply reflects an extension or acceleration of normal aging processes. Traditionally, the dissociation between fronto-striatal and temporal lobe atrophy has been proposed as reflecting different underlying processes in aging and AD (Buckner, 2004; Ohnishi et al., 2001). Many studies also showed that the temporal lobes are preferentially affected by age (Fjell et al., 2009; Pfefferbaum et al., 2013; Raz et al., 2010), even when focussing only on older adults at very low risk of AD (Fjell et al., 2013). In the current study, we showed that the medial prefrontal networks are relatively specific to aging, and already show substantial atrophy by the age at which it is likely that persons develop AD dementia. However, GM volume in most of the other morphometric networks changed almost linearly from young to old adulthood and was accelerated with AD dementia, resulting in an additive effect of both phenomena across most of the cortex. In fact, our results suggest that even the most AD-related regions are probably confounded by a strong influence of aging. These findings emphasize that by the time an individual develops sporadic dementia, the effect of age on brain atrophy that has spanned over decades is quantitatively similar, or even greater, to the effect of AD neurodegeneration.

GM volume in the temporal lobe was the best network to dissociate older adults from AD, but it was not specific to the disease. Only increased heterogeneity in the GM pattern and the volume across networks was more specific to AD. We showed that the whole-brain pattern did change over the course of aging and AD, but while cognitively normal older adults maintained a coherent pattern, this homogeneity was lost in AD patients. These results suggest that it is not the *magnitude* of atrophy in temporal brain systems that is specific to AD, but rather the *heterogeneity* that characterizes AD. Following this idea, older individuals with a GM pattern more similar to young adults had better cognitive performance and a reduced risk of converting to MCI. Importantly, this finding was independent of the total GM volume, reinforcing the idea that assessing whole-brain GM signatutre gives information about brain integrity that is independent from atrophy. Such results accord well with the concept of brain maintenance, postulating that maintaining youth-like brain integrity is associated with “healthier” aging (Nyberg et al., 2012). It has been suggested that older adults who exhibit more youth-like functional characteristics had higher cognitive performance (Samu et al., 2017; Sun et al., 2016). Adding to this idea of functional maintenance, it is possible that structural maintenance is also an important factor of successful aging. We hypothesize that preserved GM volume in the frontal cortex more specifically might contribute to maintaining a whole-brain pattern more similar to young adults, and, in turn, better cognition. In effect, the prefrontal cortex and anterior cingulate or networks involving those regions are often related to preserved cognition in old age or even “super aging” (Arenaza-Urquijo et al., 2019; Sun et al., 2016). These different ways of exploring age and AD differences reinforce the importance of looking across the lifespan to untangle underlying processes of normal and pathological aging.

There are considerable inter-individual differences in GM volumes (Alexander-Bloch et al., 2013), and it is often assumed that such differences increase with aging, due in part to early neurodegenerative processes (Jagust, 2013). Looking at changes across the lifespan and dementia allowed us to compare directly heterogeneity in GM volume across different age and disease groups. Refuting the popular view that age is associated with increased variability, we found that GM volumes across all brain systems were as variable in young adulthood as in old adulthood. Similar findings have previously been shown when only focusing on the hippocampal volume (Lupien et al., 2007), perhaps the brain region most commonly used as a structural proxy of AD-neurodegeneration (Jack Jr et al., 2015b). More generally, it is possible that inter-individual differences influence some cross-sectional differences attributed to age- or disease-related changes. Heterogeneity in GM volume in young adults could reflect cortical endophenotypes, being present since childhood (Shaw et al., 2007). lMCI- and AD-ADNI groups showed higher GM variability than young and cognitively normal older adults, suggesting that increased variability is associated with disease stage. These results also highlight the importance to consider the vast inter-individual differences when classifying a biomarker as being normal or abnormal, without refuting that diseases increase inter-individual brain variability, at least in advanced stages.

There are important methodological aspects to consider in this study. First we defined AD as clinical AD rather that preclinical AD (Sperling et al., 2011), knowing that ~20% of our “normal” older adults have probably entered the preclinical phase of AD (Jack Jr et al., 2017). Since pathology can affect neurodegeneration in the preclinical phase of the disease (Doŕe et al., 2013; Wirth et al., 2013), the inclusion of these preclinical individuals might have slightly increased our power in detecting differences between young adults and “normal” aging and/or reduced our power in detecting differences between “normal” aging and AD. For instance, by removing individuals in preclinical AD, we expect that the dissociation between normal aging and AD dementia based on whole-brain intrinsic pattern would have been even more important since the group of cognitively normal older adults would have become more homogeneous. The multiple sites and scanners are also important confounds to consider. To minimize the effect of scanner acquisition strength, we included images acquired at 3T only. Similar to another multi-cohort study on structural covariance (DuPre and Spreng, 2017), we optimized the common GM template by averaging the template of each different group so that each group is represented equally. Our results were consistent between two groups of young adults, of cognitively normal elderly and of patients with severe cognitive impairment. Also, the main findings related to aging were validated in the mono-centric lifespan Cam-CAN study.

Overall, while atrophy occurred throughout aging and disease in an additive manner, GM volume loss was not specific to AD in any brain regions. Instead, AD compounds the effects of normal aging, but was specifically characterized by higher heterogeneity in both GM volume and whole-brain pattern signature. A more accurate understanding of the GM changes differentiating aging from AD can be uncovered when looking across the lifespan. The dissociation between GM volume and the intrinsic pattern of morphometric networks could provide new perspectives in our understanding of AD and might apply to other neurodegenerative diseases.

## 4. Acknowledgements

The authors wish to acknowledge the PREVENT-AD staff, especially Jennifer Tremblay-Mercier, Ccile Madjar, as well as the Brain Imaging Center of the Douglas Mental Health Research Institute. A full listing of the PREVENT-AD Research Group members can be found here. We would also like to acknowledge the participants of the PREVENT-AD cohort for dedicating their time and energy to helping us collect these data. Part of data collection and sharing for this project was funded by the Alzheimer’s Disease Neuroimaging Initiative (ADNI) (National Institutes of Health Grant U01 AG024904) and DOD ADNI (Department of Defense award number W81XWH-12-2-0012). ADNI is funded by the National Institute on Aging, the National Institute of Biomedical Imaging and Bioengineering, and through generous contributions from the following: AbbVie, Alzheimer’s Association; Alzheimer’s Drug Discovery Foundation; Araclon Biotech; BioClinica, Inc.; Biogen; Bristol-Myers Squibb Company; CereSpir, Inc.; Cogstate; Eisai Inc.; Elan Pharmaceuticals, Inc.; Eli Lilly and Company; EuroImmun; F. Hoffmann-La Roche Ltd and its affiliated company Genentech, Inc.; Fujirebio; GE Healthcare; IXICO Ltd.; Janssen Alzheimer Immunotherapy Research & Development, LLC.; Johnson & Johnson Pharmaceutical Research & Development LLC.; Lumosity; Lundbeck; Merck & Co., Inc.; Meso Scale Diagnostics, LLC.; NeuroRx Research; Neurotrack Technologies; Novartis Pharmaceuticals Corporation; Pfizer Inc.; Piramal Imaging; Servier; Takeda Pharmaceutical Company; and Transition Therapeutics. The Canadian Institutes of Health Research is providing funds to support ADNI clinical sites in Canada. Private sector contributions are facilitated by the Foundation for the National Institutes of Health (www.fnih.org). The grantee organization is the Northern California Institute for Research and Education, and the study is coordinated by the Alzheimer’s Therapeutic Research Institute at the University of Southern California. ADNI data are disseminated by the Laboratory for Neuro Imaging at the University of Southern California.

## 5. Author Contributions

Conceptualization, A.P.B., S.V. and E.V.P.; Methodology, A.P.B., J.W.V., R.L.J. and E.V.P.; Formal Analysis, A.P.B., J.G. and E.V.P.; Resources, P.R.N., J.P., J.C.S.B. and S.V.; Writing - Original Draft, A.P.B., S.V. and E.V.P.; Writing - Review & Editing, A.P.B, J.G., J.W.V., R.L.J., P.R.N., D.L.C., J.P., J.C.S.B., S.V. and E.V.P.; Visualization, A.P.B. and E.V.P.; Supervision, J.C.S.B., S.V. and E.V.P.; Funding Acquisition, J.P., J.C.S.B. and S.V.

## 7. Methods

### 7.1. Participants

We assembled a cross-sectional dataset from four different studies (n=1019) to include cognitively normal young adults (18-35 years old), cognitively normal older adults (55-90 years old), as well as individuals who represented the clearly symptomatic portion of the AD clinical continuum (late mild cognitive impairment [lMCI] and AD dementia, 56-88 years old) to disentangle the effect of age and AD on GM changes. Demographics of this multi-cohort dataset are detailed in Table 1. Written informed consent was obtained from all participants or their legal representatives under protocols approved by the Institutional Review Boards at all participating institutions.

Young adults came from two independent open access databases: the 1000 Functional Connectomes Project (FCP) and the Human Connectome Project (HCP). The FCP is a large-scale initiative combining resting-state and structural scans from adult participants from 33 sites worldwide (Biswal et al., 2010). We specifically used data from the 198 subjects between 18-30 years old collected at the Cambridge site ([FCP-Cambridge], PI: Buckner, R.L.). The HCP consortium of several universities provides a very large dataset of participants aged 18 to 35 (Van Essen et al., 2013). From these, we used 270 HCP individuals aged between 30 and 35 years old who were gender-matched to the PREVENT-AD cohort (see below).

Cognitively normal older individuals were selected from two independent databases: the PRe-symptomatic EValuation of Experimental or Novel Treatments for AD (PREVENT-AD) cohort and the Alzheimer’s Disease Neuroimaging Initiative (ADNI) database. PREVENT-AD enrols older adults with intact cognition who have a parent or two siblings with well-documented histories of AD-like dementia, and are therefore at increased risk of AD (Breitner et al., 2016). At enrolment, they must be at least 60 years of age, or between 55-59 if fewer than 15 years from their relative’s age of symptom onset, and must be free of major neurological and psychiatric diseases. Data from the baseline visits of 295 PREVENT-AD participants (Data Release 2.0, November 2015) was used in the present study. All MRI scans were acquired at the brain imaging centre of the Douglas Mental Health Research Institute, Montreal, Canada. Cognitive performance was assessed using the Repeatable Battery for Assessment of Neuropsychological Status (RBANS) (Randolph et al., 1998). We selected a memory task of list learning (10 words over 4 trials) and a test of executive function (coding) to investigate relationships between cognition and GM features. These tests have been shown previously to be sensitive to mild cognitive impairment related to AD (Peters et al., 2014; Villeneuve et al., 2009). Cognitive data were available from 291 participants.

ADNI is a multi-site study launched in 2003 as a public-private partnership. The primary goal of ADNI has been to test whether serial MRI, positron emission tomography, other biological markers, and clinical and neuropsychological assessment can be combined to measure the progression of MCI and early AD. For up-to-date information, see www.adni-info.org. The ADNI study is divided into different phases, and data for the present analyses came from ADNI2 only. ADNI2 baseline visits for continuing participants or initial visits for newly enrolled participants were selected. 135 cognitively normal participants (Controls-ADNI) were included in the present study. Additionally, those who converted to MCI during their subsequent follow-up visits (including visits up to ADNI3) (n=18) were identified for exploratory analyses aiming at comparing different GM features between Controls-ADNI converters and those who remained cognitively normal. As a measure of cognition, we used the Alzheimer’s Disease Assessment Scale-cognitive subscale (ADAS-Cog) (Rosen et al., 1984), where higher scores represent higher degree of cognitive impairment.

Clinically impaired participants were selected from the ADNI2 database. The present study includes 50 participants with lMCI, and 71 with AD dementia. Because we sought GM changes that distinguished cognitively normal aging from advanced pathological aging, we included individuals with severe cognitive impairment only. Thus, we did not include early MCI participants, as they represent a more intermediate stage.

### Complementary analyses: Lifespan validation cohort

One limitation of the multi-cohort dataset is that participants from different studies were pooled together, bringing effects inherent to different sites, scanners and image acquisitions. To validate some of our results, we performed similar analyses using data from the Cambridge Centre for Ageing and Neuroscience (Cam-CAN) study. The Cam-CAN study is a large lifespan monocentric cross-sectional population-based study in the UK (Taylor et al., 2017). This cohort is ideal to characterize age-related GM changes. We included 647 participants aged between 18 and 88 years old with T1-weighted structural scans, from the Cam-CAN Stage 2 repository. There were approximately 100 participants in each decade, except for the range of 80 to 88 years old, which included only 44 participants. See Table S2 for a breakdown of participants per decade.

### 7.2. MRI acquisition and processing

#### 7.2.1. Image acquisition

T1-weighted structural images were acquired at 3 Tesla for all individuals. The different MRI sequences from each study are detailed in Table S3.

#### 7.2.2. Processing of the grey matter maps

T1-weighted structural images were segmented into grey matter (GM), white matter (WM), and cerebrospinal fluid (CSF) images using Statistical Parametric Mapping (SPM12), running on MATLAB version 2012a. GM images went through Diffeomorphic Anatomical Registration through Exponentiated Lie Algebra toolbox (DARTEL) (Ashburner, 2007), in which inputs are iteratively aligned to create a group-specific template. The template underwent nonlinear registration with modulation for linear and non-linear deformations to the MNI-ICBM152 template. Those initial steps were carried out separately for each group, resulting in six group-specific templates (FCP-Cambridge, HCP, PREVENT-AD, Controls-ADNI, lMCI-ADNI, AD-ADNI). Then the six templates were themselves iteratively aligned using DARTEL to create a common template in MNI space. Importantly, this common template equally weighted each group, as an attempt to have a final template more representative of all subjects. A second registration was done on each participant’s GM map to warp it with modulation to the final common template. Lastly, GM images were smoothed with an 8mm3 isotropic Gaussian kernel.

The Cam-CAN dataset was analyzed as a separate group, but underwent similar steps. All images were segmented and underwent DARTEL to create a Cam-CAN-specific template. Every GM image was aligned to the Cam-CAN template, warped with modulation to the MNI space and smoothed.

All images underwent visual quality control after segmentation and after non-linear transformation.

### 7.3. Independent Component Analysis (ICA)

ICA is a computational method to decompose multivariate data into different components by maximizing statistical independence (Beckmann and Smith, 2004). We performed ICA on the GM maps of all individuals to derive data-driven regions of GM covariance. To apply such a method on structural data, we concatenated the modulated and smoothed GM maps to create a 4D file, which became the input for the ICA. To ensure that only GM voxels were retained for the ICA, the maps were masked with a maximum probability GM mask. This mask was generated from the group-average GM, WM, and CSF images and consists of voxels with highest probability of being GM (GM > WM > CSF). ICA was performed using the toolbox MELODIC from the FSL analysis package (Jenkinson et al., 2012) version 5.0.8.

To derive common data-driven components spanning lifespan and the AD spectrum, the ICA was performed on all subjects (n=1019). There is no clear rule as to how many components to extract from an ICA (Cole et al., 2010) and we set the output at 30 components as done in Zeighami et al. (2015). Each component was thresholded at z = 3.5 (Beckmann et al., 2009) and binarized to retain the voxels that contributed significantly to the component. These thresholded IC maps are hereafter referred to as morphometric networks. The GM volume for each of the 30 morphometric networks was then extracted for each participant for further analysis.

To examine whether morphometric networks were also present in participants with severe cognitive impairment, two ICA were fit separately on lMCI- and AD-ADNI groups. For these ICAs, participants GM maps were only registered to the original template of the lMCI- and AD-ADNI groups instead of the common template to avoid deformation bias. Thirty morphometric networks were thus derived in lMCI- and AD-ADNI groups (Figure S1) and were compared qualitatively to the networks derived across all participants.

### 7.4. Statistical analyses

#### 7.4.1. Cross validation analyses

From GM volume in the 30 morphometric networks, we aimed to identify which networks were affected most specifically by aging and by AD. We grouped the FCP-Cambridge and HCP samples together as “Young adults” (n=468), the PREVENT-AD and Controls-ADNI as “Older adults” (n=430), and the lMCI- and AD-ADNI as “Alzheimer’s dementia” (n=122). We used binary logistic regression models with ten-fold cross-validation to classify (1) Young adults vs. Older adults and (2) Older adults vs. Alzheimer’s dementia, with the average GM volume in each of the 30 networks as input. We then used receiver operating characteristic (ROC) analyses and measured the area under the curve (AUC) to assess the model performance across the collated test sets. AUC were classified as follows: excellent=0.90-1, good=0.80-0.89, fair=0.70-0.79, poor=0.60-0.69, or fail=0.50-0.59 (Safari et al., 2016).

The Cam-CAN dataset was used to validate the effect of age on GM volume. Age was entered in a voxelwise regression analyses using SPM12, including sex and total intracranial volume as nuisance variables. Results are reported with a p 0.05 family-wise error (FWE) correction.

#### 7.4.2. Whole-brain GM pattern

Next, we assessed how measures of whole-brain GM pattern were influenced by aging and AD. We derived measures of GM pattern similarity by correlating the GM volume in the 30 morphometric networks of each individual to the GM volume in the 30 brain systems of every other subject. These correlations indicate how one’s whole-brain organization is similar to every other individual. This resulted in a 1019×1019 matrix of whole-brain GM pattern between all subjects (Figure 3B).

We evaluated whether there was a coherent GM pattern within each group (intrinsic pattern). Within the different groups, we calculated the average and standard deviation of correlation coefficients of GM pattern across all individuals. We then compared difference in correlation coefficients between groups using z test statistic 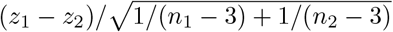 to test if the intrinsic GM pattern remained organized with aging and AD at the group level. The z test statistic formally tests if the coefficient of correlations is greater in a group compared to another given the sample size.

Next, to get a measure at the individual level, for each participant, GM volume in the 30 networks were correlated to the mean GM volume in the 30 networks of their respective group. We then used binary logistic regression and ROC analyses with 10-fold cross validation to identify whether the GM pattern within-group could differentiate Young adults from Older adults, and Older adults from Alzheimer’s dementia. This tested if whole-brain pattern homogeneity within the groups characterized aging or AD (Figure 3F). Second, to get a measure of whether the pattern itself changed with aging and AD, for each participant, GM volume in the 30 networks were correlated to the mean GM volume in the 30 networks of the Older adults group. This tested if the whole-brain pattern between groups (with older adults as the comparison point) can distinguish Young from Older adults and Older adults from AD (Figure S3).

#### 7.4.3. Heterogeneity of GM volumes

To assess group effect on GM volume across brain networks, we used repeated measures ANOVA with GM volume in the 30 networks as intra-subject measure and the six groups as the inter-subject measure. To assess variability of GM volume in aging and AD, we calculated the coefficient of variation (standard deviation/mean of GM volume in each network) in the 30 networks. We used the modified signed-likelihood ratio (MSLR) test from the R software package cvequality version 0.1.3 (Marwick and Krishnamoorthy, 2018) to test for significant differences in the coefficients of variation of GM volume between groups. A p-value smaller than 0.002 was considered significant, accounting for 30 comparisons.

To assess variability of GM volume across lifespan, coefficients of variation in the 30 networks were also calculated in the Cam-CAN dataset. The 30 networks were registered on the Cam-CAN maps and coefficients of variation in GM volume were compared across decades.

#### 7.4.4. Clinical impact of GM volume and whole-brain pattern in cognitively normal older adults

In cognitively normal older adults, we also evaluated whether GM volume or whole-brain GM pattern were related to 1) cognitive performance (PREVENT-AD and Controls-ADNI), and 2) clinical progression (Controls-ADNI only). We focused on GM volume in the network with the best discrimination between Young and Older adults (age-related network) and between Older adults and Alzheimer’s dementia (AD-related network), and on a metric representing preserved whole-brain GM pattern, i.e. pattern similarity to young adults. To test the degree to which older adults had a pattern similar/dissimilar to young adults, we correlated the GM volume in the 30 brain systems for each older adult with the mean GM volume in the 30 brains systems of the Young adults group. Correlation coefficients were Fisher z transformed.

We investigated whether the different GM features were related to cognitive performance in PREVENT-AD and Controls-ADNI groups separately using linear regression models. In PREVENT-AD, memory and executive function performance were the dependent variables and models included education and total GM as covariates. In Controls-ADNI, ADAS-Cog was the dependent variable and models included education and total GM as covariates. Analyses were run on SPSS version 20 (IBM Corp., Armonk, NY). A two-sided p-value < 0.05 was considered significant.

Finally, Mann-Whitney U tests were used to compare baseline differences in GM features between Controls-ADNI stable and converters. We also performed binary logistic regression with stable or converter status as dependent variable and GM feature as predictor, followed by ROC analyses to evaluate the discriminative accuracy of the different features. Given the small number of converters, those analyses were conducted with leave-one-out cross-validation. ROC curves were calculated across the collated test sets.

## Supplemental Information

**Figure S1:**
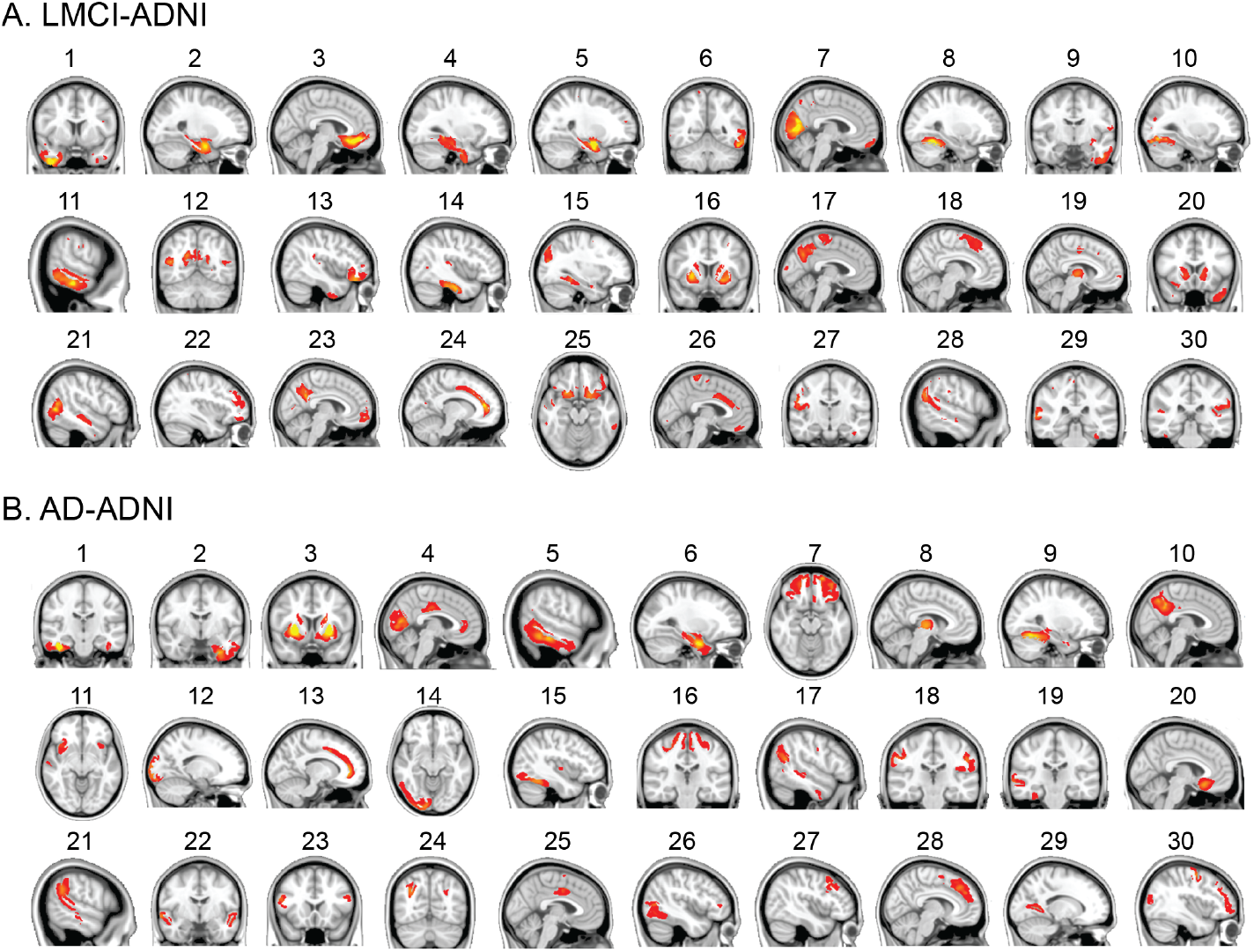
Related to Figure 2: Morphometric networks in lMCI- and AD-ADNI groups. Displays of the 30 networks derived from ICA in the lMCI-ADNI (A) and AD-ADNI (B), thresholded at Z 3.5, ordered in decreasing amount of variance explained.

**Table S1:**
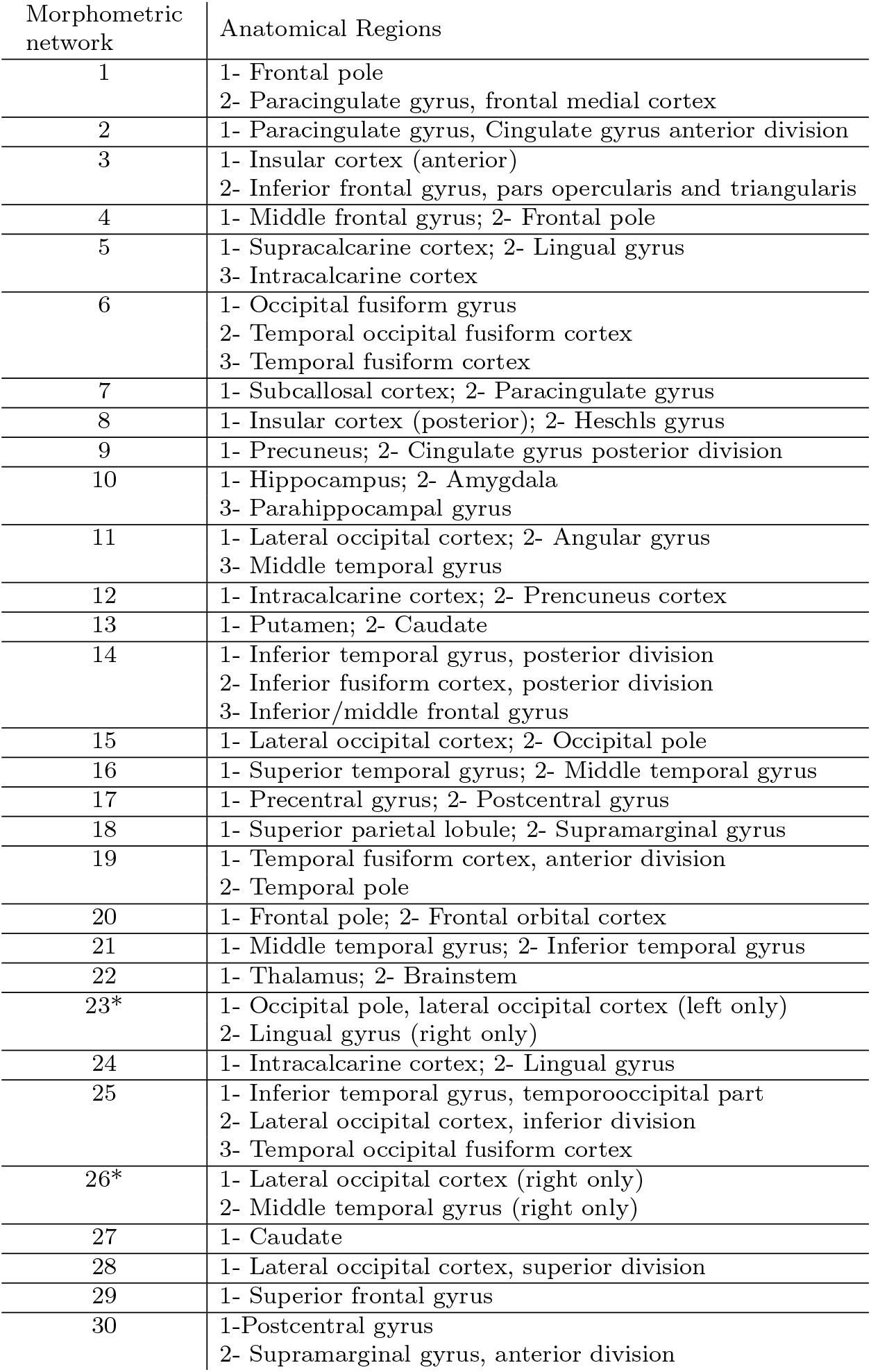
Related to Figure 2: Description of the 30 morphometric networks identified by ICA. For each network, anatomical regions highest Z-value are listed. All networks were symmetrical in the left and right hemispheres, except component 23 and 26 (marked with *). Anatomical regions were taken from the Harvard-Oxford cortical structural atlas.

| Morphometric network | Anatomical Regions |
| --- | --- |
| 1 | 1- Frontal pole<br>2- Paracingulate gyrus, frontal medial cortex |
| 2 | 1- Paracingulate gyrus, Cingulate gyrus anterior division |
| 3 | 1- Insular cortex (anterior)<br>2- Inferior frontal gyrus, pars opercularis and triangularis |
| 4 | 1- Middle frontal gyrus; 2- Frontal pole |
| 5 | 1- Supracalcarine cortex; 2- Lingual gyrus<br>3- Intracalcarine cortex |
| 6 | 1- Occipital fusiform gyrus<br>2- Temporal occipital fusiform cortex<br>3- Temporal fusiform cortex |
| 7 | 1- Subcallosal cortex; 2- Paracingulate gyrus |
| 8 | 1- Insular cortex (posterior); 2- Heschls gyrus |
| 9 | 1- Precuneus; 2- Cingulate gyrus posterior division |
| 10 | 1- Hippocampus; 2- Amygdala<br>3- Parahippocampal gyrus |
| 11 | 1- Lateral occipital cortex; 2- Angular gyrus<br>3- Middle temporal gyrus |
| 12 | 1- Intracalcarine cortex; 2- Precuneus cortex |
| 13 | 1- Putamen; 2- Caudate |
| 14 | 1- Inferior temporal gyrus, posterior division<br>2- Inferior fusiform cortex, posterior division<br>3- Inferior/middle frontal gyrus |
| 15 | 1- Lateral occipital cortex; 2- Occipital pole |
| 16 | 1- Superior temporal gyrus; 2- Middle temporal gyrus |
| 17 | 1- Precentral gyrus; 2- Postcentral gyrus |
| 18 | 1- Superior parietal lobule; 2- Supramarginal gyrus |
| 19 | 1- Temporal fusiform cortex, anterior division<br>2- Temporal pole |
| 20 | 1- Frontal pole; 2- Frontal orbital cortex |
| 21 | 1- Middle temporal gyrus; 2- Inferior temporal gyrus |
| 22 | 1- Thalamus; 2- Brainstem |
| 23* | 1- Occipital pole, lateral occipital cortex (left only)<br>2- Lingual gyrus (right only) |
| 24 | 1- Intracalcarine cortex; 2- Lingual gyrus |
| 25 | 1- Inferior temporal gyrus, temporooccipital part<br>2- Lateral occipital cortex, inferior division<br>3- Temporal occipital fusiform cortex |
| 26* | 1- Lateral occipital cortex (right only)<br>2- Middle temporal gyrus (right only) |
| 27 | 1- Caudate |
| 28 | 1- Lateral occipital cortex, superior division |
| 29 | 1- Superior frontal gyrus |
| 30 | 1- Postcentral gyrus<br>2- Supramarginal gyrus, anterior division |

**Figure S2:**
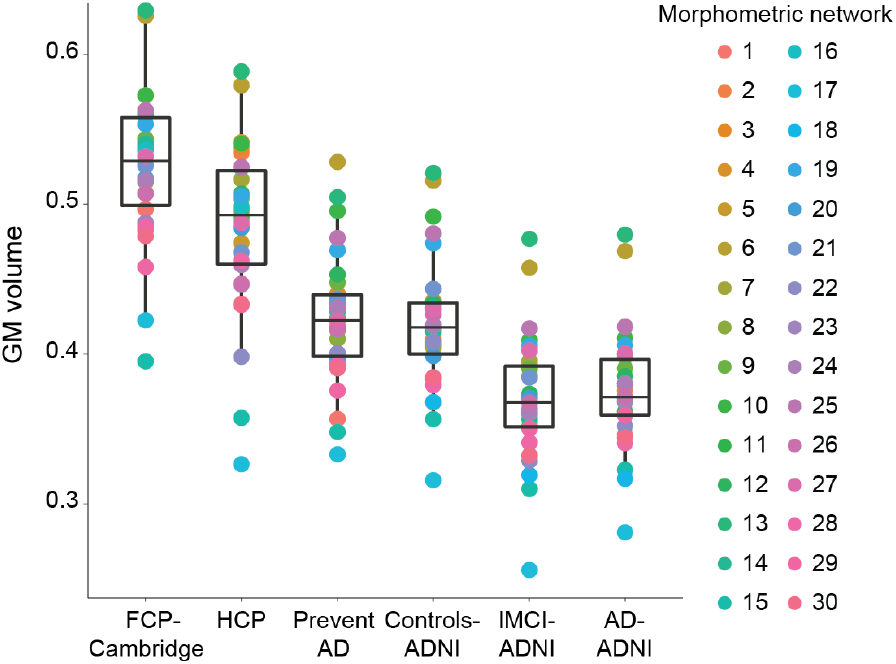
Related to Figure 2: GM volume across all morphometric networks in the multi-cohort dataset. A repeated measure ANOVA, using GM volume in the 30 networks as a repeated measure, revealed a significant group effect (F5,1014=229, p ¡ 0.001). Bonferroni post-hoc tests revealed that the FCP-Cambridge group (age range = 18-30 years) had more GM volume that the HPC group (age range = 30-35 years) (p ¡ 0.001). The two groups of young adults had more GM volume than all other groups (all ps ¡ 0.001). The PREVENT-AD and Controls-ADNI groups had GM volume similar to one another (p=1.0), but more GM volume compared to the lMCI- and AD-ADNI groups (all ps ¡ 0.001), while the latter did not differ from one another (p=1.0). Such results suggest a potential effect of age/AD or site.

**Figure S3:**
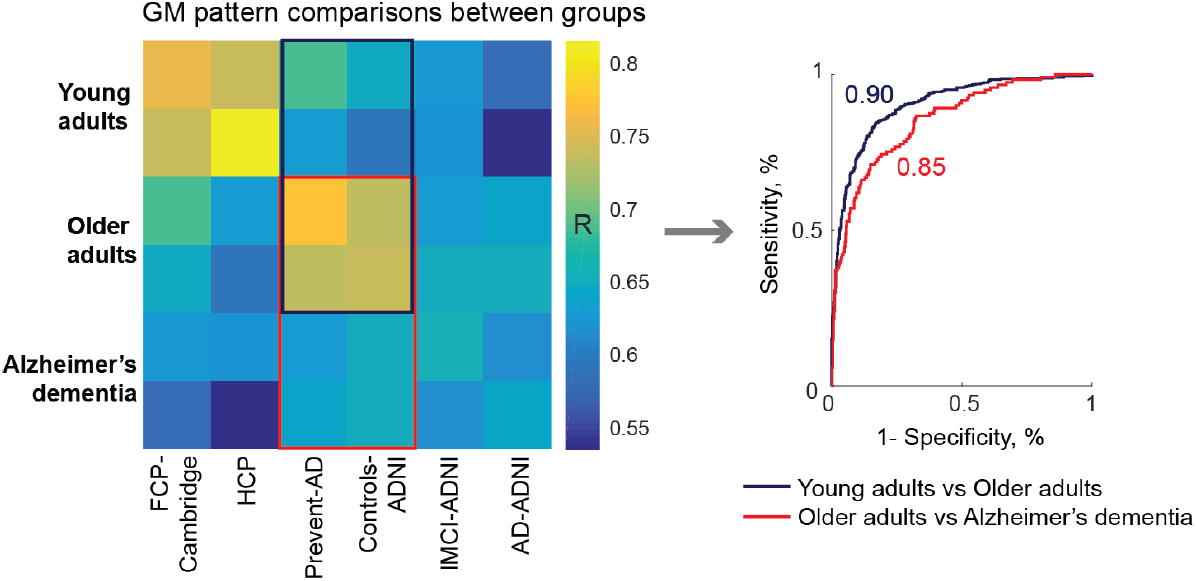
Related to Figure 3: Grey matter pattern between groups. The average GM pattern within- and between-groups are shown on the matrix (same as in Figure 5C). To get a measure at the individual level, each participants GM volumes were correlated to the mean GM volumes of Older adults. The GM pattern differed between Young and Older adults (AUC=0.90), and between Older adults and Alzheimers dementia (AUC=0.85). Overall, the GM pattern changed in youth and old adulthood and with Alzheimers dementia.

**Table S2:**
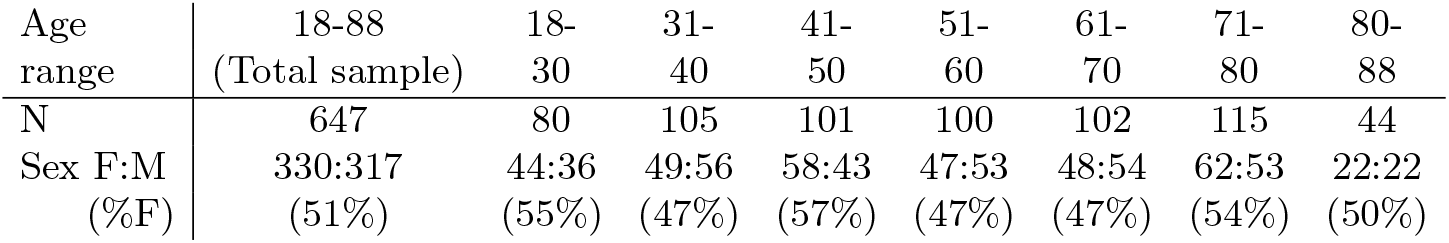
Related to Figure 4: Cam-CAN Demographics.

**Table S3:** Related to image acquisition methods: T1-weighted image acquisition parameters by study.

|  | Repetition time (ms) | Echo time (ms) | Flip angle (degree) | Field of view (mm) | Voxel size (mm) |
| --- | --- | --- | --- | --- | --- |
| FCP-Cambridge | N/A | N/A | N/A | 172x172 | $1.2 \times 1.2 \times 1.2$ |
| HCP | 2400 | 2.14 | 8 | 224x224 | 0.7x0.7x0.7 |
| PREVENT-AD, ADNI | 2300 | 2.98 | 9 | 256x256 | 1.0x1.0x1.0 |
| Cam-CAN | 2250 | 2.99 | 9 | 256x240 | 1.0x1.0x1.0 |

